# Visual Strategies During a Cooperative Mechanically Coupled Bilateral Task

**DOI:** 10.64898/2026.01.31.703066

**Authors:** Ryan T. Burgardt, Rachel L. Hawe

## Abstract

A subset of bilateral tasks requires one arm to perform a stabilizing role while the other completes a movement, such as slicing a loaf of bread. Visual attention during bilateral tasks has previously been examined with bilateral reaching tasks, demonstrating that visual attention switches between the two target locations. The goal of this study was to characterize visual attention during a cooperative mechanically coupled bilateral “stabilizing and reach” task to determine how visual attention is divided between the two limbs when one limb plays a stabilizing role. Twenty-six healthy young adults completed a robotic task in which the hands were coupled with a haptic spring. Participants were instructed to keep one hand stationary in a target while they reached for a target with the other hand, thus stretching the spring and applying a force to both arms. We found that individuals primarily fixated their gaze on the reaching target, only fixating on the stabilizing target for approximately 10% of the reaching time. Longer fixations on the reaching target were associated with faster reaching times, while longer fixations on the stabilizing target were associated with slower reaching times. While the performance of the stabilizing hand differed between the dominant and non-dominant limbs, visual strategies did not vary based on which hand was stabilizing. These results demonstrate that unlike bilateral reaching tasks in which the eyes frequently saccade between the two targets, visual guidance is primarily used for the reaching hand while minimal overt visual attention is directed to the stabilizing hand.

## INTRODUCTION

The majority of activities we perform require coordination between two limbs (Kilbreath and Heard 2005; Haaland et al. 2012; Bailey et al. 2015). A specific category of bilateral tasks is referred to as cooperative tasks, in which the hands must work together to achieve a goal (Kantak et al. 2017). In many cooperative tasks, one hand performs a stabilizing role, while the other hand performs a manipulating or reaching role. Examples of such tasks include opening a bottle, slicing a loaf of bread, or stabilizing a piece of paper while writing on it. In such tasks, especially as the hands are spaced further apart, visual attention cannot be directed to both hands at once, thus competing for visual attention (Miller and Smyth 2012; Blinch et al. 2018; Sardar et al. 2023).

In unilateral tasks, gaze fixation on the target location typically precedes limb movement and facilitates accuracy (Land et al. 1999; Mennie et al. 2007; Land 2009). Bilateral tasks present a challenge to visual strategies as visual attention must be switched between the two sides (Castiello 1996; Baldauf and Deubel 2008; Bruyn and Mason 2009; Srinivasan and Martin 2010; Blinch et al. 2018; Sardar et al. 2023). Vision during bilateral tasks has mostly been examined during bilateral aiming or reach to grasp tasks (Honda 1982; Riek et al. 2003; Bruyn and Mason 2009; Srinivasan and Martin 2010; Sardar et al. 2023). The most common strategy is to fixate on one target first, and then sequentially switch to the other target. When one target is more challenging due to either a greater distance or smaller target, there is a bias to saccade to the more challenging target first (Honda 1982; Bruyn and Mason 2009; Sardar et al. 2023). Prior work has been contradictory as to the bias in which side is fixated first when the targets are equivalent sizes and distances, with some denoting a gaze preference toward the dominant side (Honda 1982; Foerster et al. 2011; Sardar et al. 2023), others to the non-dominant side (Riek et al. 2003; Srinivasan and Martin 2010), or even no bias at all (Bruyn and Mason 2009). Additionally, while it is most common to saccade between the targets so that each target is fixated on at some point in the movement, individuals can perform bilateral reaching movements without ever fixating on either of the targets (Honda 1982).

Visual strategies may help to explain motor behavior of the limbs during bilateral tasks. For instance, in independent bilateral reaching tasks (each arm is reaching a separate target), several studies have found a “hover phase” in which one arm reaches the target first and “hovers” while waiting for the second arm to reach the target (Riek et al. 2003; Srinivasan and Martin 2010; Miller and Smyth 2012). This may be due to switching of visual attention between the two targets. The asynchrony between the two limbs in bilateral reaching tasks varies based on the spacing of the start or end target positions, likely due to greater distances between targets requiring more overt visual attention shifts and thus increased asynchrony between the limbs (Miller and Smyth 2012). Additionally, prior work has demonstrated that the required kinematics can be identical between tasks but when the goals are conceptualized differently, performance can be altered (Mechsner et al. 2001; Diedrichsen et al. 2006; Shea et al. 2016). Specifically, performance on cooperative reaching in which individuals move both limbs to control a common target is better than independent reaching (Diedrichsen 2007; Shea et al. 2016; Kantak et al. 2016). One reason for this may be that the single midpoint visual target negates the need to shift visual attention.

Visual strategies for tasks in which one arm is stationary in a stabilizing role while the other limb moves have not been examined. These tasks differ from independent bilateral reaching as only one hand is moving, however, the other must stay stationary while a load is applied to it by the moving limb. Such tasks may have particular interest for rehabilitation following a stroke, when individuals may use the more affected limb as a “helper” in the stabilizing role. Using gaze tracking while individuals perform a cooperative stabilizing and reaching task can uncover the visual strategies that are used when the limbs have disparate roles, and how the visual strategies may relate to the movement kinematics.

Stabilizing tasks reveal specific features of the specialization of each hemisphere. The dynamic dominance hypothesis posits that the dominant hemisphere is optimized for trajectory control while the non-dominant hemisphere is specialized in impedance control or limb stabilization (Sainburg 2002; Schaefer et al. 2009). Cooperative stabilize and reaching tasks using a mechanical spring between the arms have previously shown that the non-dominant arm is better at stabilizing than the dominant (Woytowicz et al. 2018). As visual attention may explain motor behavior, a question that arises is whether the visual strategies used differ between when the dominant or non-dominant arm is stabilizing. Visual strategies may also differ between hands based on a previously described specialization of the right (dominant) arm for visual feedback and the left (non-dominant) for proprioceptive feedback (Goble and Brown 2008).

The aim of this study was to use robotics and gaze tracking to characterize visual attention during a mechanically coupled task in which one hand is instructed to stay stationary while the other hand reaches to a target, stretching a haptic spring between the hands. Our objectives were to characterize the visual strategies used by healthy young adults performing this task and determine how difference in visual attention correlated with performance of both the stabilizing and reaching limbs. Additionally, we wanted to determine if visual strategies are different for when each arm plays the stabilizing role. We hypothesized that while individuals would fixate on the reaching arm more than the stabilizing arm, they would intermitted saccade to fixate on the stabilizing arm to confirm it was within the target. Additionally, we expected the fixation time on the stabilizing hand to be lower when stabilizing with the non-dominant arm, due to the non-dominant limb specialization for both stabilization and use or proprioceptive feedback.

## METHODS

### Participants

We recruited healthy young adult participants between the ages of 18 and 35 years with no known history of neurological impairment or orthopedic condition impacting their upper extremities. Twenty-six participants (mean age ± standard deviation: 25.2 ± 4.5 years; 8 males/18 female, 23 right-handed/3 left-handed) completed the study protocol. Handedness was defined as the hand used for writing. The Institutional Review Board of the University of Minnesota approved the study, and participants gave written informed consent prior to participation. While the focus of this study is on the stabilize and reach task, participants completed several different robotic tasks in a random order in the same visit which are beyond the scope of this study.

### Kinarm Exoskeleton

Evaluation of the participants utilized a Kinarm exoskeleton robot (Kinarm, Kingston, ON, Canada) with an integrated EyeLink 1000 Remote eye tracking system (SR Research, Ottawa, ON, Canada). Participants sat with their arms supported on a horizontal plane by bilateral exoskeleton robots as seen in Figure 1. The exoskeletons were adjusted to align with the participant’s shoulder and elbow joints. A solid black panel below the display prevented vision of the participant’s own arms. A virtual reality display projected their hand position feedback as white circular cursors and task related visuals (targets, spring visualization) in the same plane as their arms. The gaze tracking system was calibrated using five target locations so the coordinates of the gaze point of regard would be in the same coordinate frame as the limb positions and task visuals.

**Figure 1.**
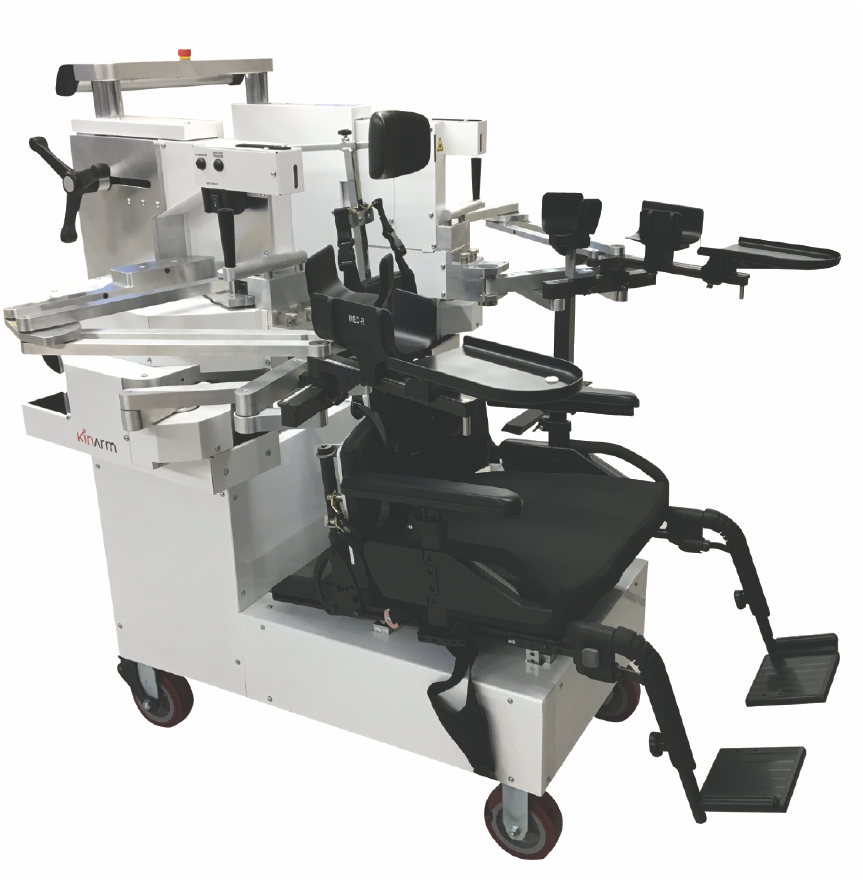
Robotic Exoskeleton Set-up. The bilateral Kinarm exoskeleton robot with adjustable arm troughs that support the arms in the horizontal plane.

### Stabilize and Reach Task

We custom programmed a task for use with the Kinarm to require participants to stretch a haptic spring between their hands to reach for a target with one hand while keeping the other hand stationary. This replicates bilateral actions where one arm serves as a stabilizing hand and the other a manipulating hand. A similar paradigm has previously been done using a mechanical spring and was adapted for this Kinarm task (Woytowicz et al. 2018; Jayasinghe et al. 2021).

The participants first reached for two red starting targets located 12 cm from the midline of the screen, and 24 cm apart from each other. The participant then held the starting target positions for a random time interval between 1500 ms and 2500 ms to prevent anticipation of the go signal. A green target then appeared on the screen in one of three positions, as shown in Figure 2A. The haptic spring forces as well as visual representation turned on when the green target appeared on the first trial and remained on until completion of all trials. Participants were instructed to stabilize on the red starting target with one hand while reaching for the green end target as seen in Figure 2B. A spring constant of 40.0 N/m was selected after piloting so the task could also be performed by participants with weakness due to neurological conditions (not included in this study). The distance and angles of the reaching targets are shown in Figure 2A. To ensure the spring force was constant across all targets, the distance from the stabilizing target to end reaching targets (the length of the spring when stretched) was kept constant (34 cm) across trials, which did vary the distance from the reaching start to end targets slightly (10 cm vs. 12.5 cm). The participant held their hand at the reaching target for 1000 ms until the targets turned off, completing the trial. After the first trial, which was considered a practice trial and not included in analysis, participants completed five blocks of trials with each block consisting of each target location in a random order, for a total of 15 trials. Prior to beginning the task, participants saw a video demonstration and received verbal instructions to stretch the spring to reach the end target as quickly as possible while keeping their other hand stationary in the target.

**Figure 2.**
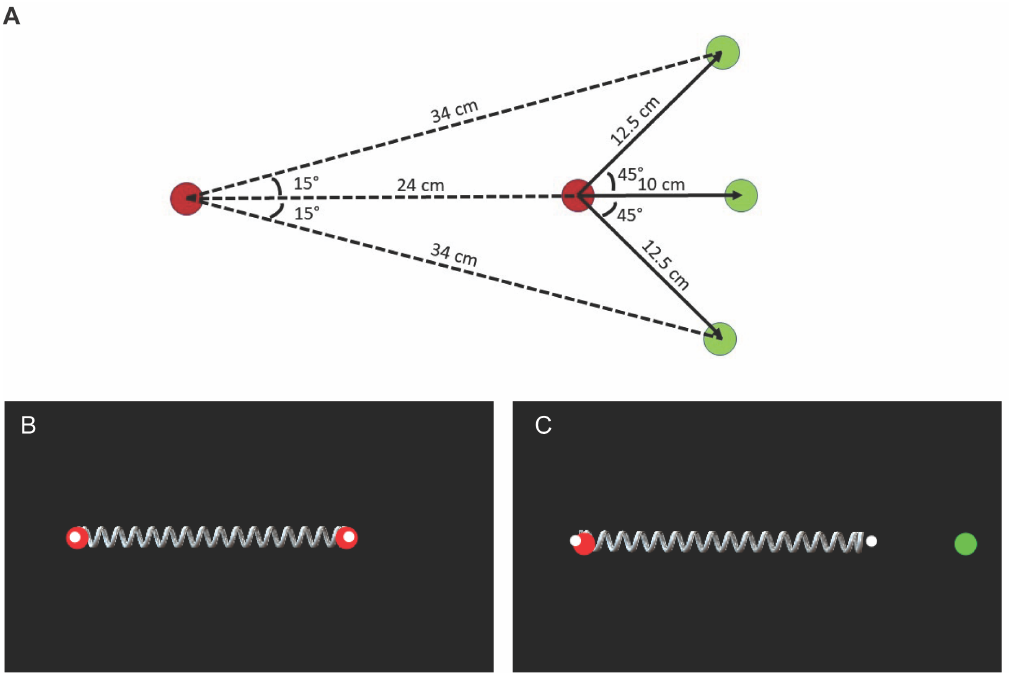
Stabilize and reach task. A) The locations and distances between targets is shown. The red circles are the start targets, and the green targets are the locations of the different end targets). The configuration is shown for when participants stabilized with the left hand while reached with the right hand. B and C show the participant’s workspace. B shows starting position is shown with the white circles providing feedback on the location of the hands. The hands begin in the red starting circles, with the spring visualized between them and providing force on both limbs. C) The position during mid-reach is shown, where the participant attempts to keep the left hand stationary on the red circle while stretching the spring to reach the green target with the right hand.

Participants completed the protocol twice, once with the dominant limb stabilizing and non-dominant limb reaching, and once with the non-dominant limb stabilizing and dominant limb reaching. Limb kinematic data was collected from the Kinarm with a sampling rate of 1000 Hz and gaze data was collecting with a sampling rate of 500 Hz.

### Limb Kinematic Analysis

Limb kinematic data from the Kinarm was low-pass filtered using a double-pass Butterworth filter with frequency cutoff at 10 Hz. Primary outcomes for the stabilizing hand were peak hand distance traveled and peak acceleration, as have been previously used in a similar paradigm (Jayasinghe et al. 2021). Peak hand distance was defined as the maximum distance away from the participant’s initial stabilizing hand position. Peak hand acceleration was defined as the maximum acceleration of the stabilizing hand. We also examined the number of trials with errors, determined by whether the stabilizing hand left the visual target. Performance of the reaching hand was measured using the movement time, calculated as from when the end target turned on until they had reached the end target.

### Gaze Analysis

Gaze data was analyzed from when the end targets turned on until movement offset of the reaching hand (Coderre et al. 2010). This analysis period was used to include the initial time period after the reaching hand first reaches the end target, as this is when individuals may look at the stabilizing hand. First, blinks and artifacts were removed and missing data was interpolated. Trials were excluded if more than 15% of the gaze data was missing, however, the limb kinematic data was still included in analysis. The gaze data was then filtered with a Savitzky Golay filter with a low-pass cutoff frequency of 20 Hz. Previously established methods that have integrated gaze tracking with the horizontal Kinarm display were then applied to classify saccades and fixations (Singh et al. 2016), which were then manually verified. We calculated the foveal visual circle to indicate what region of the 2D horizontal plane was in the participant’s central vision as a function of gaze point-of regard, distance from the eye, and calibration error (Singh et al. 2016). If the foveal visual circle overlapped with a target during a fixation, the target was assumed to be fixated. We could then determine the time each target (stabilizing and reaching) was fixated on. Primary outcome measures for gaze were the percentage of an analysis period fixated on the stabilizing target and the percentage fixated on the reaching targets. We also visualized the gaze strategies by normalizing the reaching time (0 to 100% reach completion) across all trials to determine the percentage of all trials in which each target (start, end, and stabilizing) was fixated on as the reach progressed.

### Statistical Analysis

First, to guide our statistical analysis, we visually examined trends in our outcomes across trials to determine if there was a learning effect. This confirmed that the first trial (practice) showed worse performance which may be due to the spring being initiated in this trial (it remains on for remaining 15 trials) or a learning effect. No learning effects were evident in the remaining trials. Therefore, trials 2-16 were included in the analysis. To examine if the non-dominant hand was better at stabilizing, we used paired t-tests to compare the frequency of errors (hand leaving the stabilizing target) and mean differences for dominant vs. non-dominant hands for peak hand distance traveled and peak acceleration. We also used paired t-tests to compare the percentage of time the stabilizing and reaching targets were fixated on between when the dominant and non-dominant limbs were stabilizing. Lastly, we correlated limb kinematics (movement time of reaching limb, maximum acceleration and peak displacement of the stabilizing hand) with the percent of time fixated on the reaching and stabilizing targets using repeated measures correlations (Bakdash and Marusich 2017).

## RESULTS

### Gaze Strategies

Out of 780 trials, data from gaze was acceptable and analyzed for 671 trials. Exemplar data for a participant’s limb movements and gaze fixations are shown in Figure 3. The participant initially fixated on the reaching hand’s start target (purple shaded region), followed by a longer fixation on the reaching end target (red shaded region) during most of the reaching movement which includes the main velocity peak of the reaching hand. The participant then fixated on the stabilizing hand target (blue shaded region) during the last portion of the trial, which is accompanied by small movements of the stabilizing hand.

**Figure 3:**
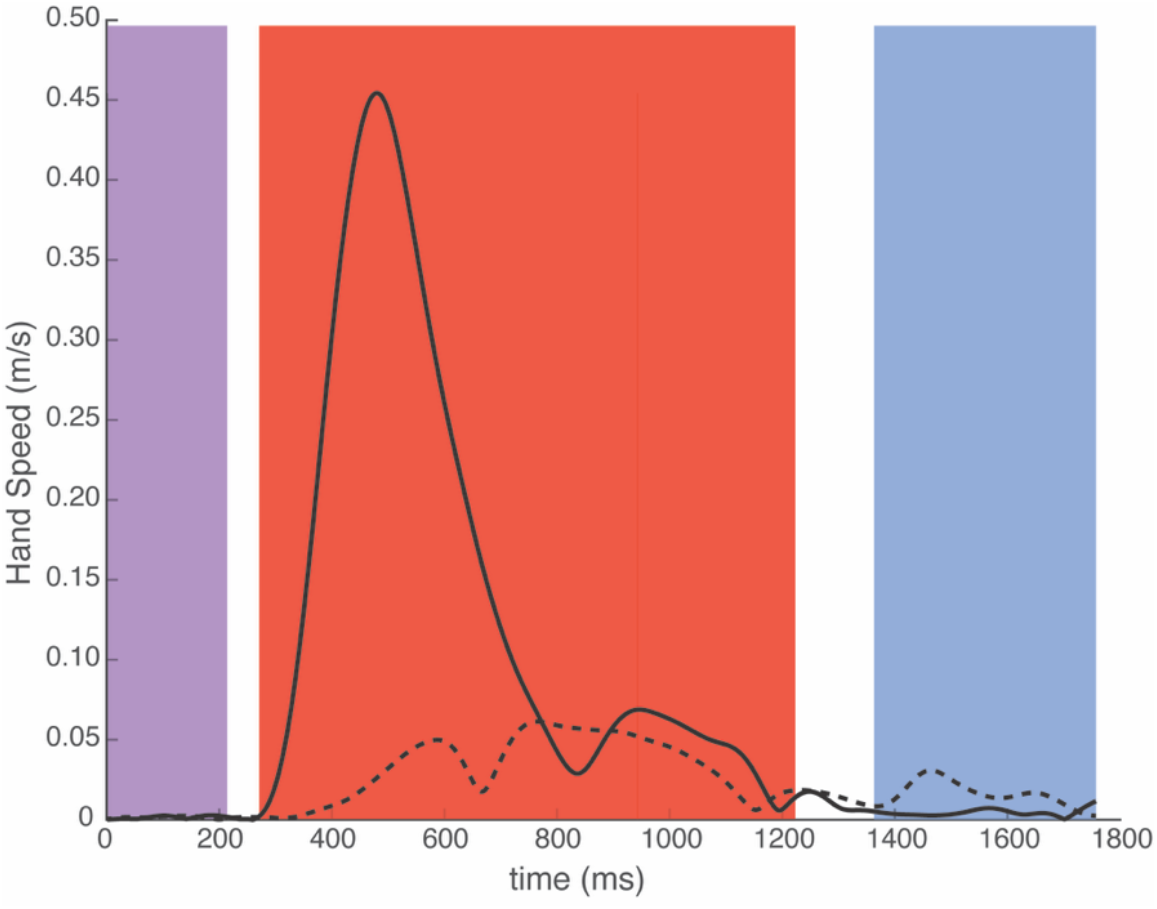
Exemplar limb and gaze fixation data. The hand speeds of the two hands (solid line is reaching hand, dashed line is stabilizing hand) are shown from when the end target appeared until movement offset following reaching the end target. The shaded regions indicate where the gaze was fixated. The purple indicates on the reaching hand’s start target, the red indicates on the reaching target, and the blue indicates on the stabilizing target.

While this was a commonly observed strategy, the gaze behavior across all trials and participants can be examined in Figure 4 which demonstrates the percentage of all trials in which the reaching start target, reaching end target, and stabilizing target were fixated on as the reach progressed from the end target turning on until movement offset.

**Figure 4:**
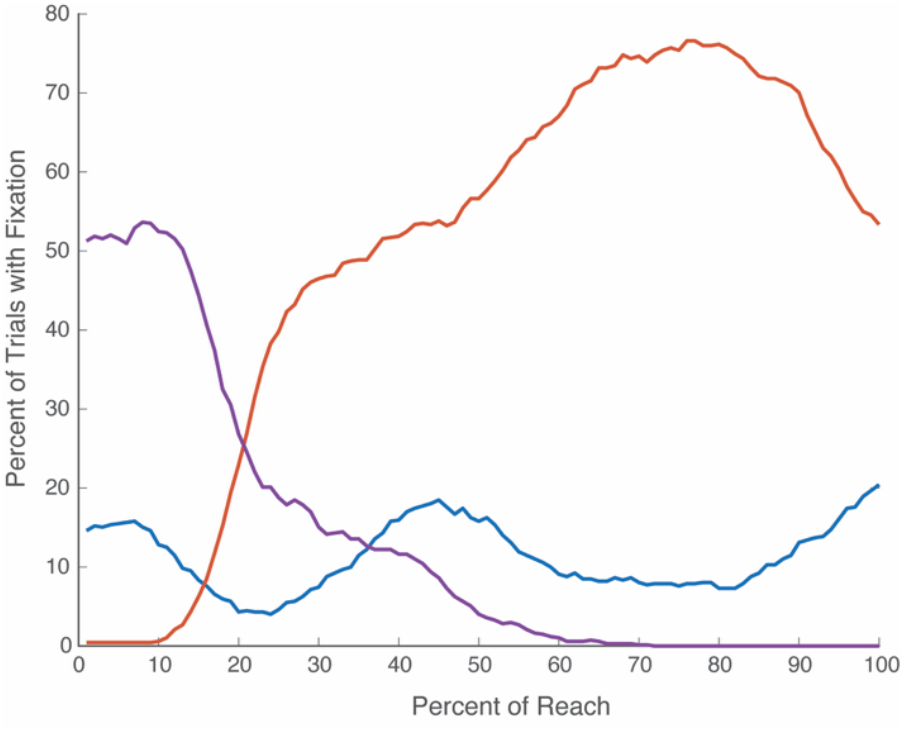
Compilation of gaze strategies across all trials. Time was normalized to examine the gaze behavior as the reach progressed, with 0 being when the end target turned on and 100% of reach being the movement offset after the end target was reached. The percentage of all trials in which the gaze is fixated on the reaching hand’s start target (purple), reaching end target (red), or stabilizing target (blue) is shown as the reach progresses.

A similar strategy to the exemplar data can be seen across the compiled data of 671 trials in Figure 4. When the end target first turns on, participants are most likely fixated on the reaching hand’s start target (>50% of trials) or the stabilizing target (∼15% of trials), the current locations of their two hands. As the reach progresses, the frequency of fixating on the end target increases, reaching a peak of nearly 80% of trials at 80% reach completion. The frequency of fixating on the end target then decreases, though still occurring in over 50% of trials, as the frequency of fixating on the stabilizing target increases in the last 20% of the reach. There is also an increase in the frequency of fixating on the stabilizing target prior to 50% of reach completion.

Notably, the task could be performed without fixating on either target. In 39% of trials, participants did not have any periods of fixation on the stabilizing target. In 6% of trials, participants did not have any fixations on the reaching target.

### Differences between dominant and non-dominant limbs

We examined differences in limb performance and gaze strategies between when the dominant and non-dominant limbs performed the reaching versus stabilizing role. For trials with the dominant hand stabilizing, the hand left the stabilizing target on 27% of trials, and for trials with the non-dominant hand stabilizing, the hand left the stabilizing target on 22% of trials. There was a significant difference in the number of trials with errors between dominant and non-dominant hands (p=0.046). The performance of the dominant versus non-dominant hand as stabilizing hand is shown in Figure 5 for the measures of peak distance traveled and peak acceleration. For peak distance traveled, the non-dominant hand traveled significantly less than the dominant hand (dominant: 0.86 cm ± 0.25 cm, non-dominant: 0.74 cm ± 0.19 cm; p=0.032). The peak acceleration of the stabilizing hand did not significantly differ between the dominant and non-dominant hands (dominant: 0.615 ± 0.308 m/s^2^, non-dominant: 0.555 ± 0.246; p=0.068).

**Figure 5.**
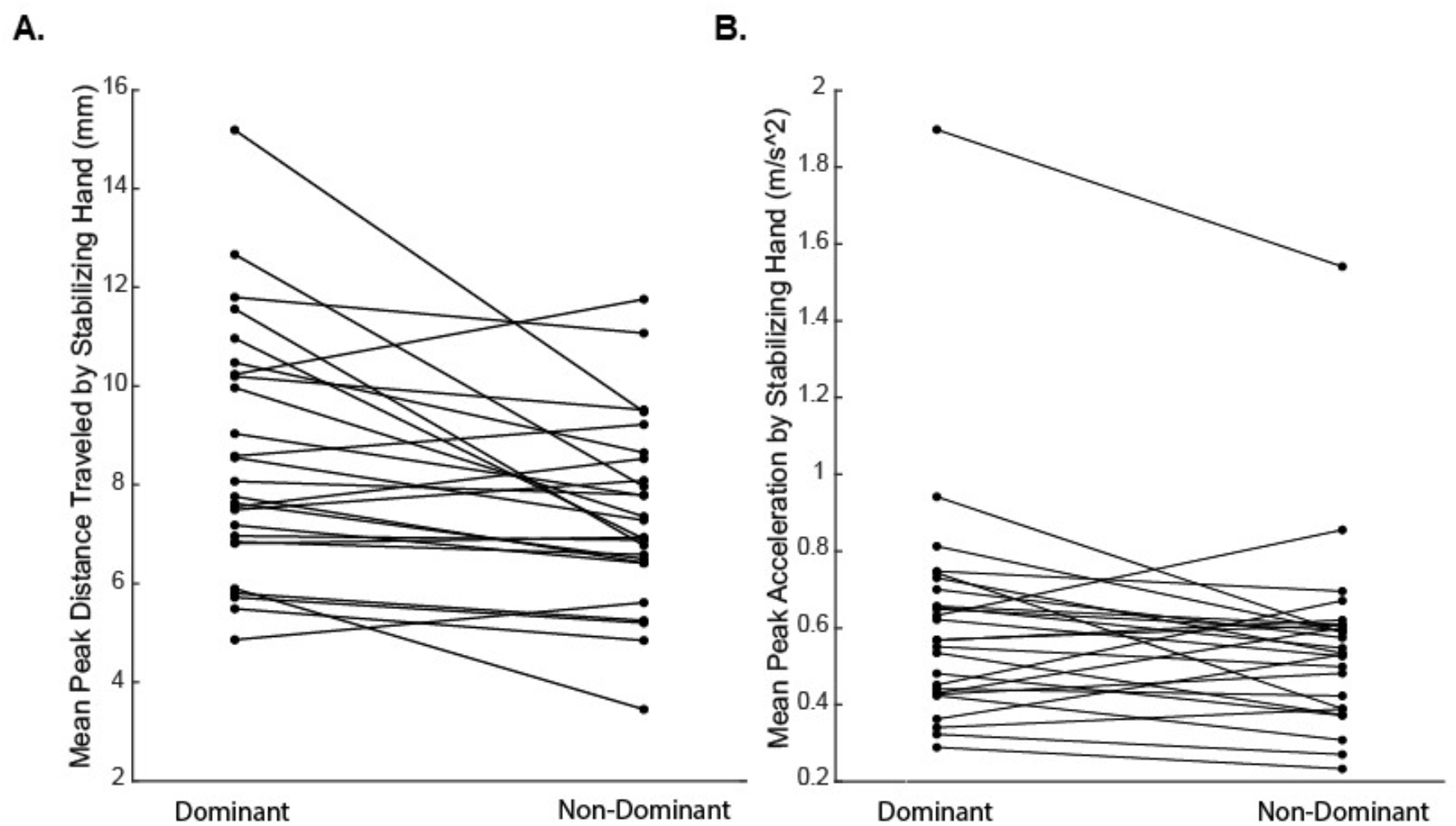
Kinematic Measures of Stabilizing Hand Performance in Dominant versus Non Dominant Limbs. Each point represents A) a participant’s mean (across trials) peak distance traveled by the stabilizing hand and B) a participant’s mean peak acceleration by the stabilizing hand. A line connects the means of dominant and non-dominant hands for each participant. The peak distance traveled was significant higher for the dominant hand than non-dominant hand, no significant difference was found between hands for peak acceleration.

We then examined if there are differences in the gaze strategies between when the dominant versus non-dominant limb is stabilizing. Figure 6 shows the percentage of the reach in which the stabilizing target (A) and reaching end target (B) were fixated by each participant for trials with their dominant and non-dominant limbs stabilizing.

**Figure 6:**
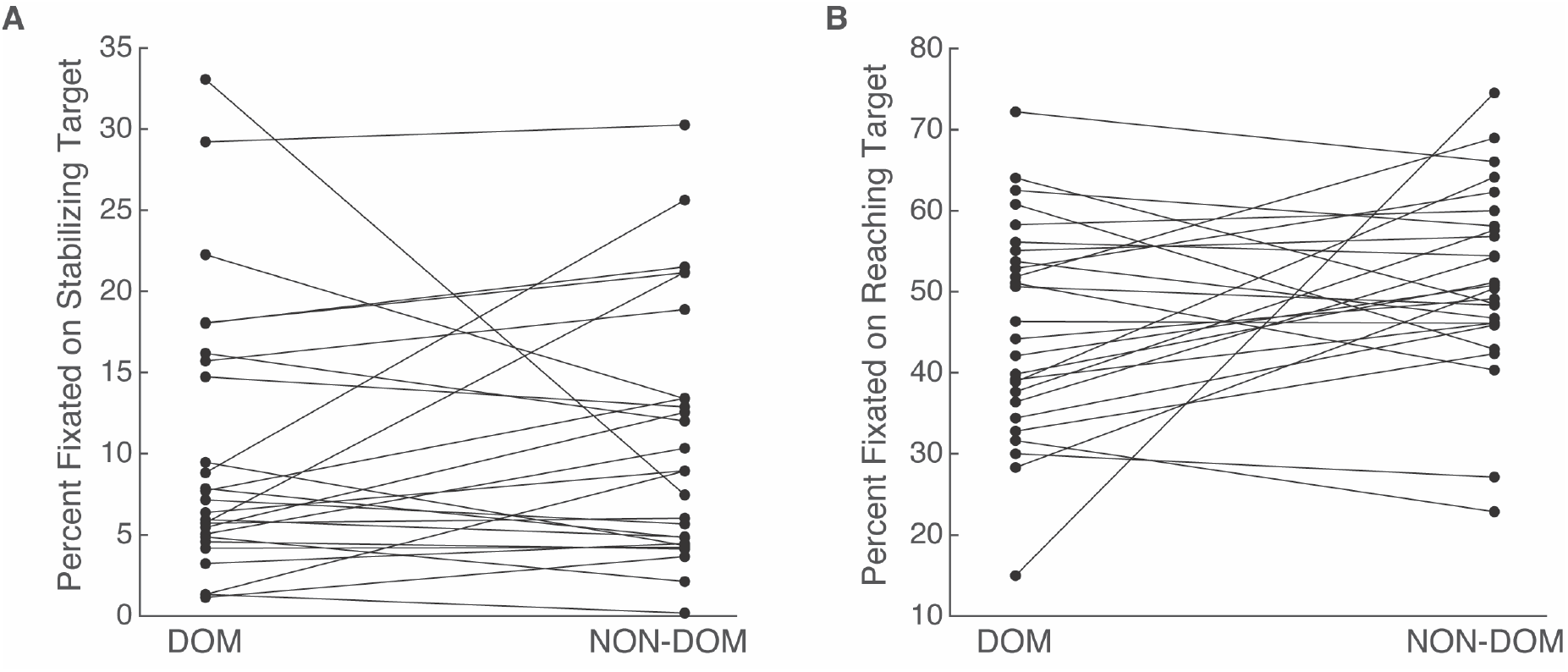
Fixations on stabilizing and reaching targets for dominant versus non-dominant limbs. A) The percentage of the trial (from end targets turning on until movement offset) that the gaze was fixated on the stabilizing target is shown with each point representing the average across trials for a participant for when the dominant (DOM) or non-dominant (NON-DOM) hand was stabilizing. B) The percentage of the trial that the gaze was fixated on the reaching target is shown with each point representing the average across trials for a participant for when the dominant (DOM) or non-dominant (NON-DOM) hand was reaching. No significant differences in fixations were found for either target between limbs.

For stabilizing, the mean percentage of the reach in which the stabilizing target was fixated on was 10.1±8.4% for the dominant arm as stabilizing arm, and 10.9±7.9% for the non-dominant arm as stabilizing arm. There was no difference in fixation on the stabilizing target between limbs (p=0.622). For fixations on the reaching target, the mean percentage of the reach in which the reaching target was fixated on was 45.6±12.2% for the dominant hand as reaching hand, and 51.4±11.6% for the non-dominant hand as reaching hand. There was no significant difference in fixations on the reaching target between limbs (p=0.075).

### Relationship between Gaze and Limb Kinematics

Figure 7 depicts the percentage of each reach that the gaze was fixated on the reaching or stabilizing target with the movement time of the reaching limb, across all trials for all participants. A repeated measures correlation was used to correlate gaze and hand speed data across all trials. We found a negative relationship between the percent of the reaching period that gaze was on the reaching target and the movement time (p<0.001, r=-0.206), and a moderate correlation between percent of time the gaze was fixated on the stabilizing target and reaching time (r=0.429, p<0.001). Correlations between the percent fixation on the stabilizing target and maximum acceleration and peak displacement of the stabilizing hand were not significant (p=0.056 and p=0.225, respectively), nor were correlations between percent time fixated on the reaching target and peak displacement or maximum acceleration of the stabilizing hand (p=0.412 and p=0.195, respectively).

**Figure 7.**
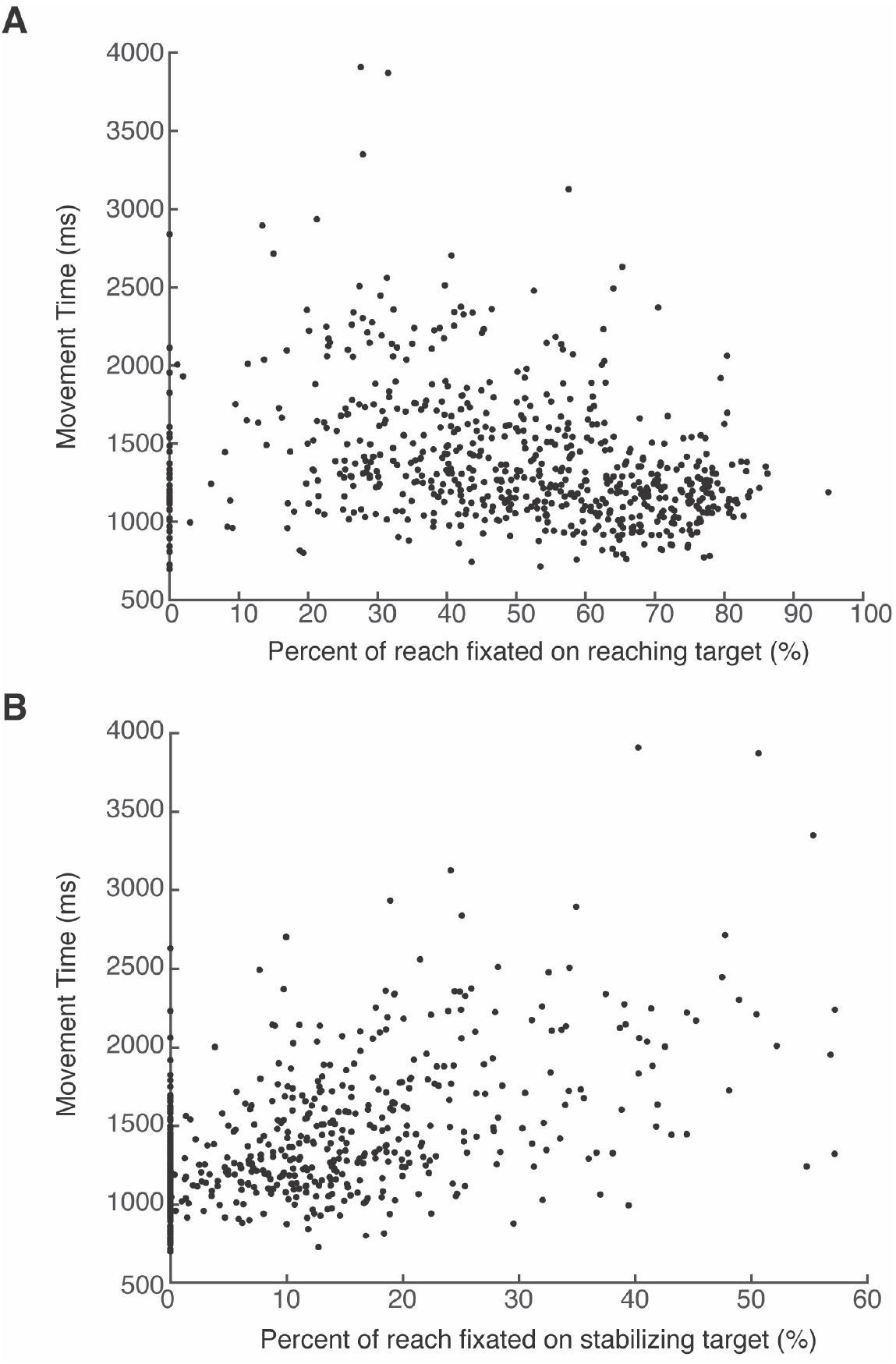
Relationship Between Visual Attention and Hand Kinematics. Each figure shows correlations of mean reaching hand speed and percentage of an analysis period fixated on A) the reaching target and B) the stabilizing target. All trials from all participants are represented as single points.

## DISCUSSION

The objective of this study was the characterize gaze strategies during a mechanically coupled cooperative bilateral task in which one arm had a stabilizing role and the other arm reached for a target, thus stretching the spring between the hands. We found that gaze was primarily directed at the reaching target. Participants were most likely to fixate on the stabilizing target just prior to movement offset, likely to confirm the stable position. There was no difference between the percentage of time the targets were fixated when it was the dominant versus non-dominant hand in each role. We found that fixations on the reaching target were correlated with faster movement times of the reaching hand, and fixations on the stabilizing target were correlated with slower movement times of the reaching hand. This work provides the first description of visual strategies during a mechanically coupled bilateral task in which the hands are playing district roles.

Examining gaze strategies can give insight into overt visual attention. Many bilateral tasks, including the one used in this study, present the challenge of dividing visual attention between two targets. Previous work has shown that individuals will primarily fixate on one target until the hand has reached or nearly reached the target before saccading to the other target, thus focusing visual attention on one goal at a time until it is achieved (Honda 1982; Riek et al. 2003; Bruyn and Mason 2009; Srinivasan and Martin 2010; Sardar et al. 2023). However, in these previous studies, both hands have been moving toward targets, whereas here, one hand is stationary. We found that as with prior work on bilateral reaching, gaze during the reach was primarily directed toward one target, in this case that of the reaching hand, until the end of the reach, when visual attention may switch to the other target. However, compared to prior work, the other hand’s target (stabilizing target) was fixated to a lesser degree, accounting for approximately 10% of the reaching time and absent in 39% of trials. Fixation on the stabilizing target was most common just prior to movement offset. This demonstrates that visual attention is primarily used to plan the reaching movement and ensure accuracy of the reaching hand, which was the primary focus until the end of the movement when individuals may “check” the position of the stabilizing hand. Overall, the stabilizing hand was largely not under direct visual guidance, which differs from visual strategies for tasks in which both hands are moving. We can also explain differences in the fixation on the reaching versus stabilizing hand based on the level of difficulty of each hand. In previous examinations of bilateral reaching, vision is primarily directed to the more challenging target, typically the smaller or further target (Honda 1982; Bruyn and Mason 2009; Sardar et al. 2023). This is presumed to be that higher accuracy demands require greater visual guidance. In our protocol, the targets were of the same size, however, only one required movement, and therefore could be viewed as the more challenging target.

We found that foveal vision was not used to monitor the stabilizing limb but rather used to confirm the position once the reaching hand had completed or nearly completed its movement. While this is the first investigation of visual strategies for stabilizing in a bilateral task, we can compare our results to what is more broadly known about the role of vision for stabilization. Prior work has examined limb stabilization during perturbation tasks (Bourke et al. 2015). Our protocol differs from this work as the “perturbation” is being provided by the spring being stretched by the other hand, thus it is not unexpected but rather self-generated. Additionally, individuals have proprioceptive information on the “perturbation” from both hands, as the spring force is applied to both. These factors likely decrease the visual reliance needed for stabilization compared to prior work with unexpected perturbations. Work in both controls and stroke patients showed that endpoint errors were smaller when visual feedback was provided, however, the rest of the correction to the perturbation did not change with visual feedback (Bourke et al. 2015). In our work, the use of vision at the end of the trial could be used to correct end point error.

Our results are consistent with the established roles of central and peripheral vision for limb movements. Central vision is needed to plan accurate reaching movements, which is consistent with the immediate fixation on the reaching target once the target turns on (González-Alvarez et al. 2007). Both central and peripheral vision can perform online error correction, however, central vision is more effective (Lawrence et al. 2006; González-Alvarez et al. 2007). While the stabilizing target was rarely fixated on with central vision, we can assume that peripheral vision was used to monitor the stabilizing hand’s position during the reach. As gaze tracking does not measure peripheral vision, future work would be needed to either manipulate individual’s field of view or force fixation locations to further examine the role of peripheral vision for a stabilize and reach cooperative task.

When examining how the gaze strategies impact motor performance of the limbs, we found that longer fixations on the reaching target were weakly associated with faster reaches, while longer fixations on the stabilizing target were moderately associated with slower reaches. This shows that directing visual attention to the reaching target may benefit the movement (faster reaches), however, this is not as strong an effect as the negative impact that focusing visual attention on the stabilizing limb has on the reaching limb. The weak correlation between fixation on the reaching target and movement time is likely due to both the fact that visual strategies only play one role in behavior, as well as the ability to perform visually guided movements without directly fixating on the target. Individuals may have fixated in the vicinity of the reaching target but not directly on the target, which would not have been measured as time fixated on the target but still may be beneficial to the control of the reach. The effect on speed is in line with prior work showing that when only peripheral vision is available (i.e. not fixating on the target), movements are slower and less direct (Sivak and MacKenzie 1990; King et al. 2010). Prior studies in bilateral reaching have found that in a subset of trials, individuals can successfully reach to a target that is never fixated on (Honda 1982; Srinivasan and Martin 2010), supporting the notion that fixation on the target is not required. However, when individuals do fixate on the stabilizing target, this directs visual attention away from the reaching arm, thus having a stronger negative impact on the reaching limb. No significant correlations were found between the gaze behavior and the stabilizing limb. This may suggest that the stabilizing limb performs this task mostly in the absence of overt visual attention or central vision. When examining the strategies, the stabilizing arm is most looked at towards the end of the trial, more as confirmation rather than stabilizing under visual control. This indicates that central vision is needed more for the reaching movement than the stabilizing movement, which may instead be performed with co-contraction and feedback from peripheral vision and proprioception.

One of the objectives of the study was to examine if there are differences in gaze strategies based on which hand performs the stabilizing role, as the hands are specialized for both different feedback (visual versus proprioceptive) and movement goals (dynamic dominance hypothesis). While we found differences between the dominant and nondominant sides in the limb kinematics on one of the two measures of the stabilizing limb, we did not find differences in gaze behavior based on which limb was performing the stabilizing role. Prior work has shown that the dominant arm is under more visual guidance while the non-dominant is specialized for proprioceptive feedback (Goble and Brown 2008). However, these feedback specializations have not been examined in a cooperative bilateral task before. The haptic spring renders the hands to be mechanically coupled and working cooperatively, and increases proprioceptive feedback between them. This may decrease the laterality effects in visual versus proprioceptive feedback previously found. Previous studies on bilateral reaching have shown conflicting results on whether gaze is preferentially directed toward the dominant or non-dominant limb, which likely suggests laterality is specific to the task protocol. We do acknowledge that our findings of differences in limb kinematics were less robust than have been previously found (Woytowicz et al. 2018; Jayasinghe et al. 2021), likely due to a lower spring force that was selected to reduce fatigue. Future work could determine if visual strategies change based on the strength of the spring.

While this study was in neurologically intact individuals, there are implications for individuals with stroke or other neurologic conditions. For individuals who have had a stroke, their more affected limb is likely to play a stabilizing role in bilateral tasks, sometimes referred to as being used as a “helper” hand. Our results on healthy young adults show that gaze is primarily directed toward the reaching target. Future work is needed to determine if this same strategy would be used in individuals with stroke stabilizing with their more affected limb. Impairments in proprioception may require them to direct more visual resources to stabilize the limb, however, our results show that increased fixation on the stabilizing target decreases the speed of the reaching limb. This could mean that an altered visual strategy in stroke would impair the function of the less affected limb, causing functional limitations. Additionally, prior work has shown that vision cannot be effectively used in individuals with stroke to correct for proprioceptive impairments (Semrau et al. 2018). In the future we will extend this protocol to individuals with stroke to determine if they do exhibit altered visual strategies compared to controls, as well as if their proprioceptive abilities relate to visual strategies.

There are several limitations to the current study. First, participants completed a limited number of trials (16 with each hand, with the first being excluded as practice), and some trials were excluded due to poor gaze quality. However, this is in line with prior studies examining gaze strategies in bilateral coordination, where trials ranged from 3 per condition in just 6 participants (Srinivasan and Martin 2010) to 10 per condition in 15-19 participants (Bruyn and Mason 2009; Sardar et al. 2023). The limited number of trials was due to developing a protocol that would be feasible in individuals with stroke who have greater levels of fatigue. This task was also part of a larger battery of assessments outside the scope of this study. Poor gaze quality causing excluded trials is a challenge with gaze tracking on the horizontal plane. While our setup has the benefit of allowing for the tracking of gaze in the same plane as the limbs, gaze artifacts are more likely to occur with proximal targets as the upper eye lid lowers. There is also a potential limitation that the force exerted by the spring was too low and did not challenge the stabilizing arm enough. In pilot testing, we examined many different spring strengths and ultimately erred on relatively light resistance to reduce fatigue and be feasible when this protocol is applied to neurologic populations. The low level of resistance may be why we did not find an effect of side in all measures for the limb kinematics as has been found in previous studies (Woytowicz et al. 2018; Jayasinghe et al. 2021). When considering functional tasks that require a stabilizing component, the resistance is highly variable, ranging from lightly stabilizing paper while writing and slicing a loaf of bread, to opening a challenging jar. Future work would be required to determine how gaze strategies may change depending on the level of stabilizing force required. Lastly, the overall way this protocol may generalize to functional tasks is unknown. This task requires reaching movements rather than more dexterous and fine movements, and limits movement to the horizontal plane. The targets (and therefore hands) were also spaced to ensure that only one could be fixated at once. Our interpretation of our results should therefore not be overgeneralized but rather show how visual attention is divided between the two limb’s roles. In stabilizing tasks where the two hands work in proximity, visual attention may not need to be divided in the same manner.

In conclusion, our study was the first to examine overt visual attention during a mechanically coupled bilateral cooperative task in which one hands reaches and the other stabilizes. We found that vision was primarily used to plan and execute the reaching movements and only used to “check” the position of the stabilizing hand. We found that visual attention on the stabilizing target did not impact stabilizing hand performance, however negatively impacted the reach, as it directed visual attention away from the reach. Future research will extend these findings to individuals with stroke, who may demonstrate different strategies due to impaired proprioception.

## ACKNOWLEDGEMENTS

The authors acknowledge Triet Lu and Samantha Timanus for assistance with data collection. This work was funded by a Grant-in-Aid from the University of Minnesota to RLH.

## STATEMENTS AND DECLARATIONS

The authors declare that they have no conflict of interest.

## Notes

### Competing Interest Statement

The authors have declared no competing interest.

